# Egr1 is a sex-specific regulator of neuronal chromatin, synaptic plasticity, and behaviour

**DOI:** 10.1101/2023.12.20.572697

**Authors:** Devin Rocks, Eric Purisic, Eduardo F. Gallo, John M. Greally, Masako Suzuki, Marija Kundakovic

**Author notes:** Correspondence should be addressed to M.K.

## Abstract

Sex differences are found in brain structure and function across species, and across brain disorders in humans^1-3^. The major source of brain sex differences is differential secretion of steroid hormones from the gonads across the lifespan^4^. Specifically, ovarian hormones oestrogens and progesterone are known to dynamically change structure and function of the adult female brain, having a major impact on psychiatric risk^5-7^. However, due to limited molecular studies in female rodents^8^, very little is still known about molecular drivers of female-specific brain and behavioural plasticity. Here we show that overexpressing Egr1, a candidate oestrous cycle-dependent transcription factor^9^, induces sex-specific changes in ventral hippocampal neuronal chromatin, gene expression, and synaptic plasticity, along with hippocampus-dependent behaviours. Importantly, Egr1 overexpression mimics the high-oestrogenic phase of the oestrous cycle, and affects behaviours in ovarian hormone-depleted females but not in males. We demonstrate that Egr1 opens neuronal chromatin directly across the sexes, although with limited genomic overlap. Our study not only reveals the first sex-specific chromatin regulator in the brain, but also provides functional evidence that this sex-specific gene regulation drives neuronal gene expression, synaptic plasticity, and anxiety- and depression-related behaviour. Our study exemplifies an innovative sex-based approach to studying neuronal gene regulation^1^ in order to understand sex-specific synaptic and behavioural plasticity and inform novel brain disease treatments.

## Introduction

Sex-specific characteristics of the brain are dynamically shaped throughout life, as a result of the actions of gonadal hormones, sex chromosomes, and the environment^1,4^. During the reproductive period in females, cycling ovarian hormones, oestrogens and progesterone, play a critical role in brain plasticity relevant to reproductive and non-reproductive behaviours including mating, feeding^10,11^, learning^12,13^, reward processing^14,15^, and anxiety- and depression-related behaviours^6,7^. In addition, ovarian hormone shifts are associated with increased risk for neuropsychiatric conditions such as anxiety and depression disorders in women^7,16^. Thus, revealing molecular mechanisms and drivers underlying female-specific brain plasticity is of crucial importance for understanding sex-specific brain physiology and disease risk.

So far, molecular studies in neuroscience have focused on the male brain^8^, and our knowledge of sex-specific molecular mechanisms in the brain is therefore limited^1^. It has been assumed that a specific group of transcription factors, known as immediate early gene (IEG) products, drive chromatin^17^ and synaptic^18^ plasticity across sexes, yet the molecular basis of sex differences in synaptic and behavioural plasticity remains obscure. Here, we leveraged previous findings from a physiological mouse model, which indicated that Egr1 is an oestrous cycle-dependent IEG and a candidate chromatin regulator driving synaptic and behavioural changes across the oestrous cycle in females^9^. By overexpressing Egr1 in neurons of the ventral hippocampus (vHIP) of ovariectomized (OVX) female mice and in males, here we examine two fundamental questions: i) whether Egr1 acts as a sex-specific IEG and chromatin regulator in neuronal cells; ii) whether Egr1 induction is sufficient to reproduce the effects of the high-oestrogenic proestrus stage on neuronal gene regulation, synaptic plasticity, and behaviour in ovarian hormone-depleted females.

## Results

### Egr1 overexpression in vHIP neurons induces proestrus-like behaviour in female mice

We previously identified Egr1 as a putative driver of gene regulatory changes in vHIP neurons across the female mouse oestrous cycle^7^ (**Fig. 1a**). During the high-oestrogenic, proestrus phase of the cycle, we found chromatin opening surrounding Egr1 binding sites coupled with increased Egr1 gene expression, compared to the low-oestrogenic, dioestrus stage^7^. To better understand the cycle-dependent functions of Egr1, here we first determined Egr1 expression in the vHIP across all phases of the oestrous cycle (**Fig. 1b**). The rodent oestrous cycle is 4-5 days long, comprising four stages (proestrus, oestrus, metestrus, dioestrus), with cyclical oestrogen and progesterone levels (**Fig. 1a**). While Egr1 shows a complex cyclical expression pattern across the oestrous cycle (**Fig. 1b**), its initial rise coincides with the rise in oestrogen in proestrus^9^, increased expression of its target gene *Ppp1r1b* (**Extended Data Fig. 1**) and reduced anxiety-related behaviour, as evidenced by increased time spent in the open arms of the elevated plus maze (EPM, **Fig. 1c**). These findings imply that Egr1 induction and its transcriptional activity drive behavioural changes across the cycle. To determine whether Egr1 induction is sufficient to reproduce the proestrus-like behavioural state in the absence of ovarian hormones, we overexpressed Egr1 in vHIP neurons of ovariectomized (OVX) females alongside a group of age-matched males that allowed us to evaluate sex-specificity (**Figure 1d, Extended Data Fig. 2**). OVX females show similar anxiety indices to low-oestrogenic, cycling females (**Extended Data Fig. 3**). However, OVX female mice overexpressing Egr1 in vHIP neurons exhibited reduced anxiety- and depression-related behaviours including increased centre entries in the open field, increased time in the open arms of the EPM, and reduced time spent immobile in the forced swim test, relative to eGFP-overexpressing controls (eGFP, **Figure 1e**). Notably, none of these behaviours were affected in males following Egr1 overexpression (**Figure 1f**), implying a sex-specific effect. Further, Egr1 overexpression did not affect overall locomotor activity in either sex (**Extended Data Fig. 4**), indicating that its effect is specific to anxiety- and depression-related behavioural indices.

**Figure 1.**
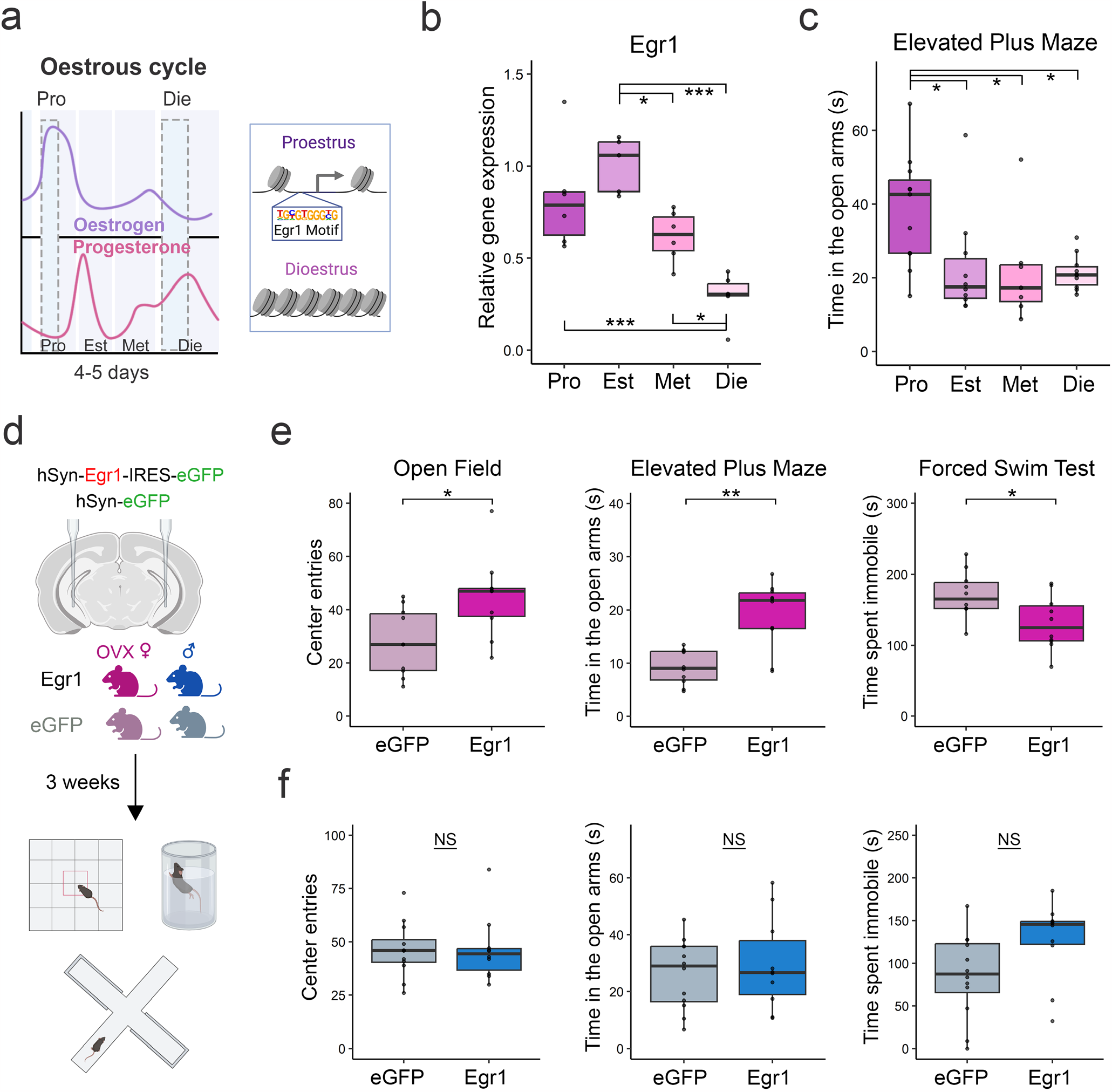
Neuronal Egr1 expression drives sex-specific behaviours. Ovarian hormone changes (**a**) drive cyclical Egr1 expression in the ventral hippocampus (n=5-6/group) (**b**) and elevated plus maze behaviour (n=8-12/group) (**c**) across the oestrous cycle in female mice; analysed with one-way ANOVA with Holm’s post hoc test. AAV-mediated Egr1 overexpression in ventral hippocampal neurons in vivo affects a battery of anxiety- and depression-related behaviours in a sex-specific manner (**d**), by mimicking the high-estrogenic proestrus phase in ovariectomized (OVX) female mice (**e**) with no effect in males (**f**), as compared to their respective eGFP controls (n=10-12/group/sex); analysed with Welch two-sample t-test. Box plots (box, 1st-3rd quartile; horizontal line, median; whiskers, 1.5xIQR); ^*^, P<0.05; ^**^, P<0.01; ^***^, P<0.001; NS, non-significant. Pro, proestrus (purple); Est, oestrus (light-purple); Met, metestrus (pink); Die, dioestrus (light-pink).

### Egr1 drives neuronal gene expression in a sex-specific manner

With Egr1’s well defined role as an IEG and a transcription factor^19^, we reasoned its sex-specific effects on behaviour could arise due to its influence on neuronal gene expression. Specifically, vHIP neurons are known to drive behaviours such as EPM^20^. Thus, we performed the same AAV-mediated Egr1 overexpression experiment in male and OVX female mice, followed by gene expression analysis (RNA-seq) on purified neuronal (NeuN+) nuclei isolated from the targeted vHIP (**Figure 2a**). Egr1 overexpression markedly altered neuronal gene expression in both sexes, with 1209 differentially expression genes (DEGs) found in females and 1680 DEGs in males (Padj<0.05, **Figure 2b**). As expected, Egr1 was one such DEG and exhibited an AAV-mediated ∼13-fold increase in expression in both sexes (**Extended Data Fig. 5a**). While we observed a similar magnitude of gene expression changes in males and females, the majority of DEGs were sex-specific, with only 37.3% (785/2104) of DEGs being shared between the sexes (**Figure 2c**). Gene set enrichment analysis (GSEA) revealed male-specific effects on the expression of genes related to peptide and neurotransmitter transport (**Figure 2d)**. By contrast, female-specific enrichment includes genes related to excitatory synapses, thyroid hormone signalling, and, importantly, behavioural fear response (**Figure 2d**), consistent with the female-specific behavioural effect we observed (**Fig. 1e-f**). To further explore the transcriptional basis for Egr1’s sex-specific behavioural effects, we examined gene sets that show group-by-sex interaction in the GSEA analysis and found enrichment for several pathways related to GABA receptor activity (**Figure 2d**), which is known to mediate anxiety-related phenotypes.^21^

**Figure 2.**
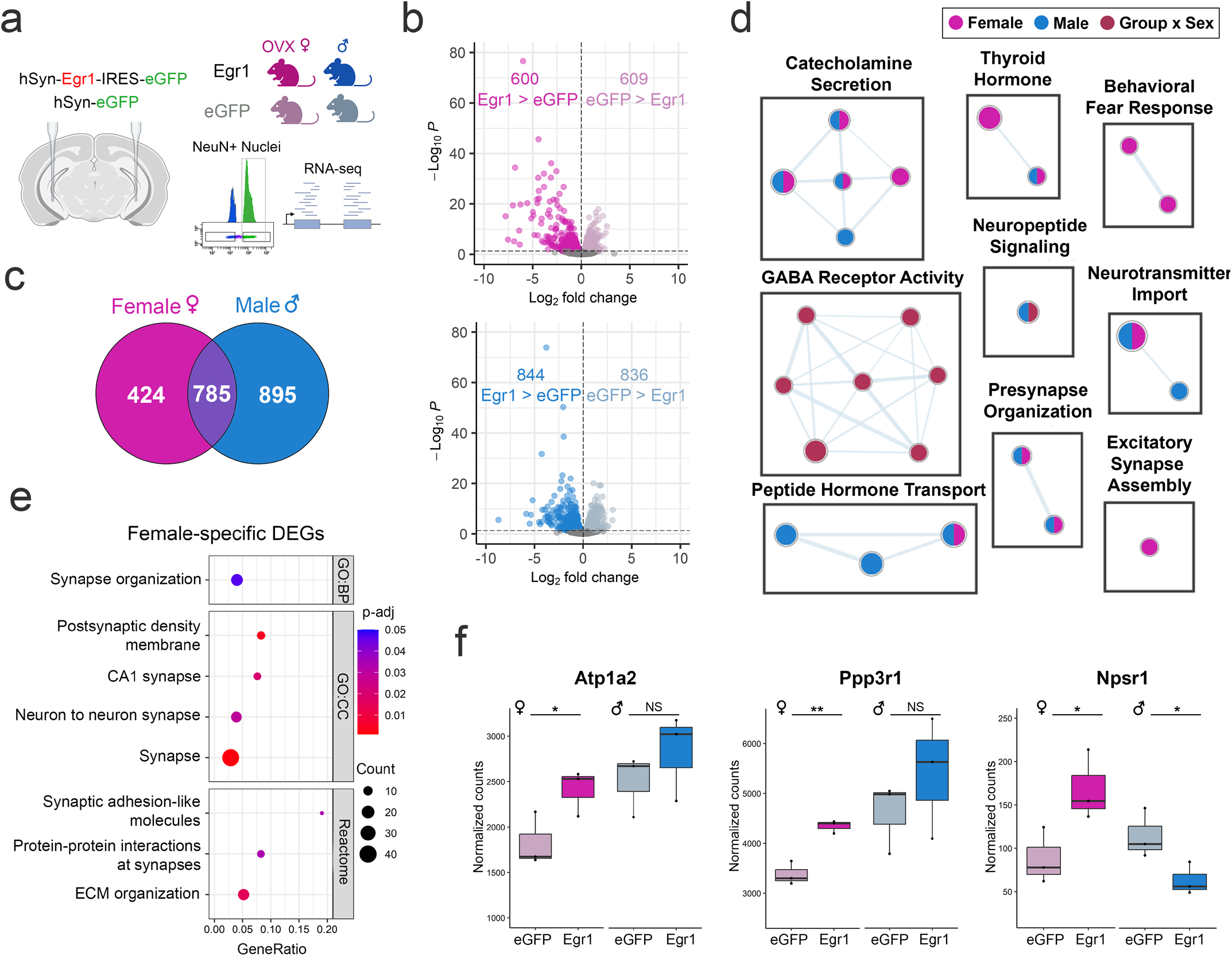
Egr1 overexpression induces sex-specific gene expression changes in vHIP neurons. **a**. RNA-sequencing was performed on neuronal (NeuN+) nuclei purified from the vHIP of males and OVX females injected with either Egr1 or eGFP AAVs. **b**. >1,000 genes show changes in gene expression in females and males (Padj<0.05, Benjamini-Hochberg correction for multiple testing); **c**. Only 37.3% (785/2104) of DEGs are shared between the sexes; **d**. GSEA analysis shows sex-specific terms including Behavioral Fear Response; Enrichment Map depicts nodes representing terms/pathways enriched in one or more gene list (denoted by node colour), while edges represent genes shared between pathways. **e**. Female-specific genes are enriched for synaptic function-related GO terms; Dot plot colours indicate p-values and dot size corresponds to the number of overlapping genes between the list and the term/pathway (Count); the x-axis corresponds to the ratio of genes in the input list to genes in the term/pathway (GeneRatio). **f**. Sex-specific DEGs include genes implicated in anxiety- and depression-related behaviours (*Atp1a2, Ppp3r1, Npsr1*). RNA-seq data for individual genes is shown using normalized count plots (n=3 biological replicates/group/sex). Box plots (box, 1st-3rd quartile; horizontal line, median; whiskers, 1.5xIQR); ^*^, P<0.05; ^**^, P<0.01; NS, non-significant; Benjamini-Hochberg correction for multiple testing.

Next, we performed a Gene Ontology (GO) enrichment analysis focused on the 424 DEGs identified exclusively in females (**Figure 2c**), and found that these genes were enriched for numerous terms related to synaptic function (**Figure 2e**), which implicates Egr1 in selectively regulating a set of synapse-related genes in females. While effects of Egr1 overexpression likely result from its cumulative impact on gene expression, we identified several anxiety-relevant example genes with expression patterns that are concordant with Egr1’s behavioural effects (**Figure 2f**). *Atp1a2*, for instance, is a gene in the Behavioral Fear Response term enriched specifically in females (**Figure 2d**). *Atp1a2* deficient-mice exhibit enhanced fear and anxiety behaviours^22^, and we found that *Atp1a2* expression in vHIP neurons is upregulated in females and unchanged in males following Egr1 overexpression (**Figure 2f**). Another example is *Ppp3r1*, encoding calcineurin B, which is a member of multiple synapse-related terms that were enriched in female-specific DEGs (**Figure 2e**). Systemic inhibition of calcineurin was found to increase anxiety- and depression-related behaviours in male mice^23^, and its expression is upregulated in females and unchanged in males after Egr1 overexpression (**Figure 2f**). Finally, *Npsr1*, encoding the neuropeptide S receptor 1, was the only gene that exhibited the opposite pattern of expression changes across sex after Egr1 overexpression (**Extended Data Fig. 5b**). *Npsr1* has been implicated in altered anxiety indices in mice^24^ and humans^25^, and, notably, its expression is upregulated in females and down-regulated in males (**Figure 2f, Extended Data Fig. 5b**) following Egr1 overexpression.

### Egr1 opens neuronal chromatin in a sex-specific manner

Given the previous work implicating Egr1 in chromatin regulation within vHIP neurons over the oestrus cycle in females^9,26^, we wondered whether altered chromatin organization is a feature of Egr1’s sex-specific gene regulatory functions. To this end, following vHIP-targeted Egr1 over-expression, we performed chromatin accessibility (ATAC-seq) analysis on vHIP neuronal (NeuN+) nuclei purified from male and OVX female mice (**Figure 3a**). Including both Egr1 and eGFP control mice, we identified a total of 205,781 accessible chromatin regions in vHIP neurons of males and 205,105 in those of females (**Figure 3b**). Of these regions, 26.5% (54,442) in males and 25.7% (52,776) in females were accessible in a group-dependent manner, indicating that Egr1 overexpression led to substantial reorganization of neuronal chromatin (**Figure 3b**). As Egr1 was identified as being a putative “opener” of chromatin^9^, and consistent with previous findings of another IEG, cFos, inducing chromatin accessibility following neuronal activity^17^, we focused our subsequent analyses on regions that gained accessibility following Egr1 overexpression (Egr1 gained-open regions). An overlap of these regions between sexes revealed remarkable sex-specificity, with only 26.6% (10,504/39,541) of gained-open regions being shared between males and females, while 42.8% (16,921) were unique to females and 30.6% (12,116) were unique to males (**Figure 3b**). We next asked whether these regions gain accessibility indirectly, downstream of Egr1’s transcriptional effects or, alternatively, if they may be targeted by Egr1 specifically, as a function of their DNA sequence. Strikingly, motif analysis revealed, in both sexes, that more than half of regions that gain accessibility after Egr1 overexpression contain the Egr1 motif (**Figure 3c, e.g. Extended Data Fig. 6**), providing evidence that Egr1 mediates opening of neuronal chromatin targeted to specific motif-containing regions. While this was previously demonstrated for cFos in the male brain^17^, our study reveals the first IEG product that acts as a sex-specific molecular driver of chromatin accessibility in neurons.

**Figure 3.**
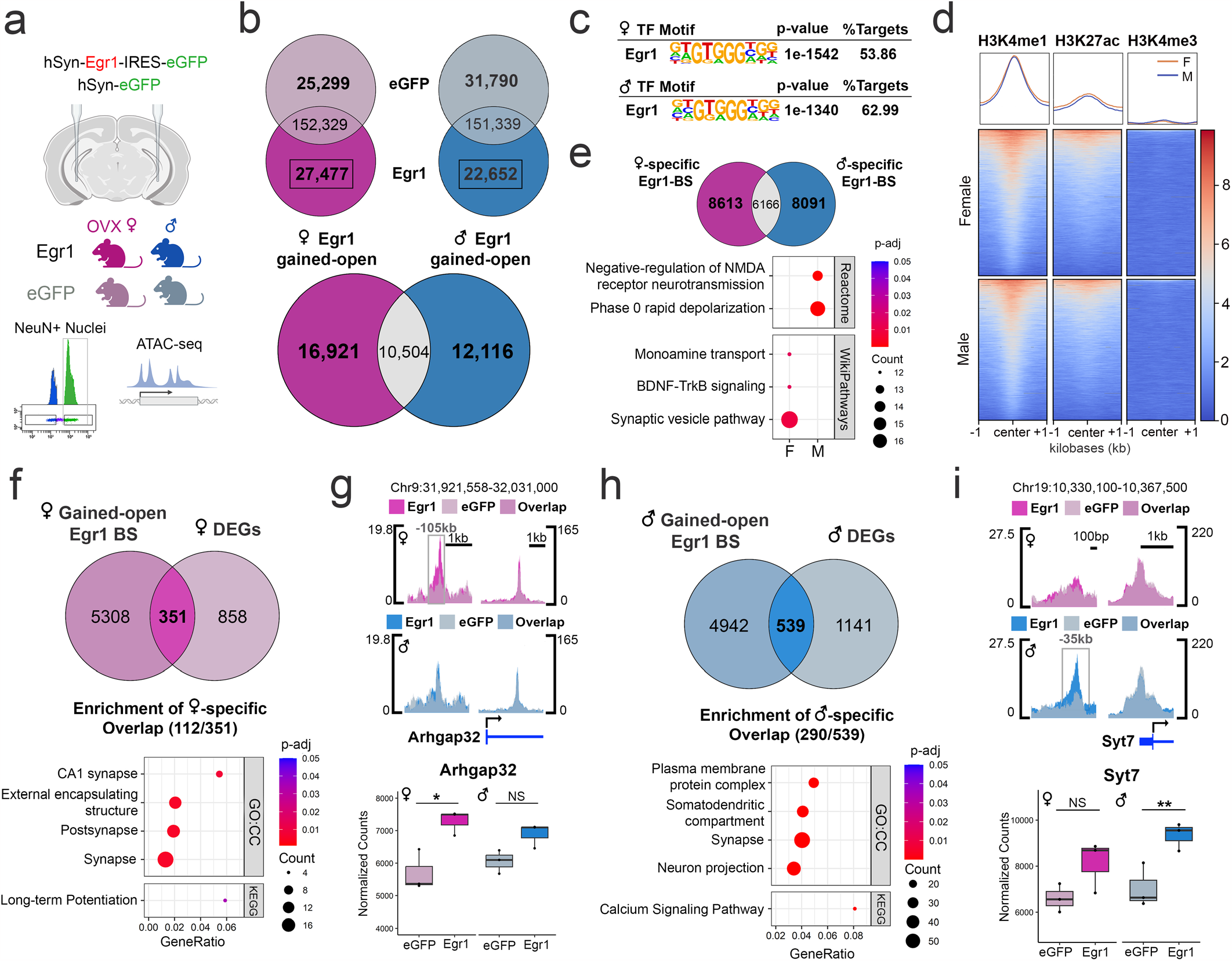
Egr1 overexpression induces sex-specific chromatin changes in vHIP neurons. **a**. ATAC-seq was performed on neuronal (NeuN+) nuclei purified from the vHIP of males and OVX females injected with either Egr1 or eGFP AAVs. **b**. Egr1 overexpression induces extensive reorganization (opening and closing) of vHIP neuronal chromatin, with only 26.6% (10,504/39,541) of gained-open regions being shared across sexes. **c**. In both males and females, Egr1-induced open regions are enriched for the Egr1 motif; Egr1 motif-containing gained-open regions: **d**. are primarily located in enhancers (overlapping with the enrichment of H3K4me1/H3K27ac but not H3K4me3 histone marks) and **e**. affect different sets of genes in males and females. **f**. In females, 29.0% (351/1209) of DEGs are associated with gained-open chromatin containing the Egr1 motif, and, of those, female-specific DEGs are enriched for synaptic function-related genes. **g**. An exemplary gene is *Arhgap32* which is associated with female-specific chromatin opening and gene expression. **h**. In males, 32.1% of all DEGs (539/1680) are associated with Egr1 motif-containing gained-open chromatin and, of those, male-specific DEGs are enriched for a different set of synaptic function-related genes. **i**. An exemplary gene is *Syt7* which is associated with male-specific chromatin opening and gene expression. Dot plots; colours indicate p-values and dot size corresponds to the number of overlapping genes between the list and the term/pathway (Count); the x-axis corresponds to the ratio of genes in the input list to genes in the term/pathway (GeneRatio). ATAC-seq data is shown using SparK plots of group-average normalized ATAC-seq reads (n = 4 biological replicates/group/sex). RNA-seq data is shown using normalized count plots (n=3 biological replicates/group/sex). Box plots (box, 1st-3rd quartile; horizontal line, median; whiskers, 1.5xIQR); ^*^, Padj<0.05; ^**^, Padj<0.01; NS, non-significant; Benjamini-Hochberg correction for multiple testing. GO:CC, Gene Ontology Cellular Component.

To explore the functional role of Egr1-induced gained-open regions containing Egr1 binding sites, we overlapped their sequences with previously published ChIP-seq data on histone modifications in hippocampal neurons^27^. Importantly, we found relative enrichment of H3K4me1 and H3K27ac, marking active enhancers^28^, and depletion of H3K4me3, marking gene promoters^29^ (**Figure 3d**), indicating that neuronal enhancers are the primary substrate for targeted opening of chromatin by Egr1. Similar to the total Egr1-induced gained-open regions, an overlap between males and females of gained-open regions with Egr1 binding sites revealed that the majority (73.0%, 16,704/22,870) are sex-specific (**Figure 3e**). Pathway analysis on genes annotated to these sex-specific regions show male-specific enrichment for pathways related to NMDA receptor signalling and rapid depolarization, and female-specific enrichment for monoamine transport-, BDNF signalling-, and synaptic vesicle-related pathways (**Figure 3e**), indicating that Egr1 initiates chromatin opening near different gene sets with distinct functions in males and females.

### Egr1-induced chromatin opening mediates gene expression changes

By overlapping ATAC-seq and RNA-seq data from the Egr1 overexpression experiments, we next explored whether Egr1’s chromatin effects were associated with effects on gene expression. A general overlap revealed that the majority of DEGs in both sexes (**Fig. 2b**), 69.7% (843/1209) in females and 73.1% (1228/1680) in males, are associated with at least one chromatin region that changes accessibility following Egr1 overexpression (**Extended Data Fig. 7**). This indicated that, in general, Egr1-induced chromatin changes mediated gene expression changes. However, we wondered which subset of these Egr1-induced DEGs were associated specifically with opening of chromatin targeted to the Egr1 motif. In females, 29.0% (351/1209) of DEGs are associated with gained-open chromatin containing the Egr1 motif (**Figure 3f**). Of these DEGs, 31.9% (112/351) are female-specific and enriched for terms and pathways related to synaptic function and long-term potentiation (**Figure 3f**). An illustrative example demonstrating Egr1’s sex-specific effects on gene regulation is *Arhgap32* (**Figure 3g**), a brain-specific Rho GTPase that has been implicated in regulating dendritic spine density^30^. Egr1 overexpression induces chromatin opening within the Egr1 motif-containing region upstream of the *Arhgap32* locus in females only, which is further associated with a female-specific increase in gene expression (**Figure 3g**).

In males, we observed a similar involvement of Egr1’s targeted chromatin opening in gene regulation, where 32.1% of all DEGs (539/1680) are associated with Egr1 motif-containing gained-open chromatin (**Figure 3h**). Of these, 53.8% (290/539) are male-specific DEGs and are enriched for terms and pathways related to synapses, neuronal projections, and calcium signalling (**Figure 3h**). An example is *Syt7*, encoding synaptotagmin 7, which plays a role in vesi-cle-mediated neurotransmitter release^31^. An Egr1 motif-containing region upstream of the transcription start site (TSS) becomes more accessible after Egr1 overexpression in males but not females, which corresponds to male-specific upregulation of *Syt7* expression (**Figure 3i**). These results demonstrate that Egr1 overexpression mediates opening of neuronal chromatin surrounding Egr1 binding sites in both sexes, with distinct genomic regions targeted in males and females associated with sex-specific effects on gene expression.

### Egr1 overexpression partially recapitulates the proestrus-associated transcriptional program in vHIP neurons

Having demonstrated that Egr1 overexpression in vHIP neurons reproduces a proestrus-like behavioural phenotype in OVX females (**Fig. 1e**), we next examined whether Egr1 overexpression similarly recapitulates molecular phenotypes associated with physiological ovarian hormone cycling in vHIP neurons. Leveraging our previously generated RNA-seq data which measured gene expression in vHIP neurons over the oestrus cycle^9^, we overlapped the identified cycle-related DEGs with DEGs resulting from Egr1 overexpression in OVX females. This analysis revealed that 18.6% (32/172) of oestrus cycle DEGs similarly undergo expression changes in OVX females overexpressing Egr1 (**Figure 4a**), a proportion of genes that is consistent with the 24.1% of oestrus cycle DEGs with chromatin changes that we previously found to have Egr1 binding motifs^9^. Two example genes which illustrate the role of Egr1-driven gene expression in oestrus cycle-dependent behavioural phenotypes are *Arid1a* and *Gatad2b. Arid1a* encodes a subunit of the SWI/SNF chromatin remodelling complex, perturbations of which have been associated with anxiety-related phenotypes^32^. Importantly, the up-regulation of *Arid1a* expression observed during the high-estrogenic, proestrus phase in cycling females is mimicked by Egr1 overexpression in OVX females (**Figure 4b**). *Gatad2b* encodes a subunit of the NuRD chromatin remodelling complex, whose levels are associated with anxiety-related behaviour^33^, and is also concordantly upregulated in proestrus females and OVX females overexpressing Egr1 (**Figure 4b**).

**Figure 4.**
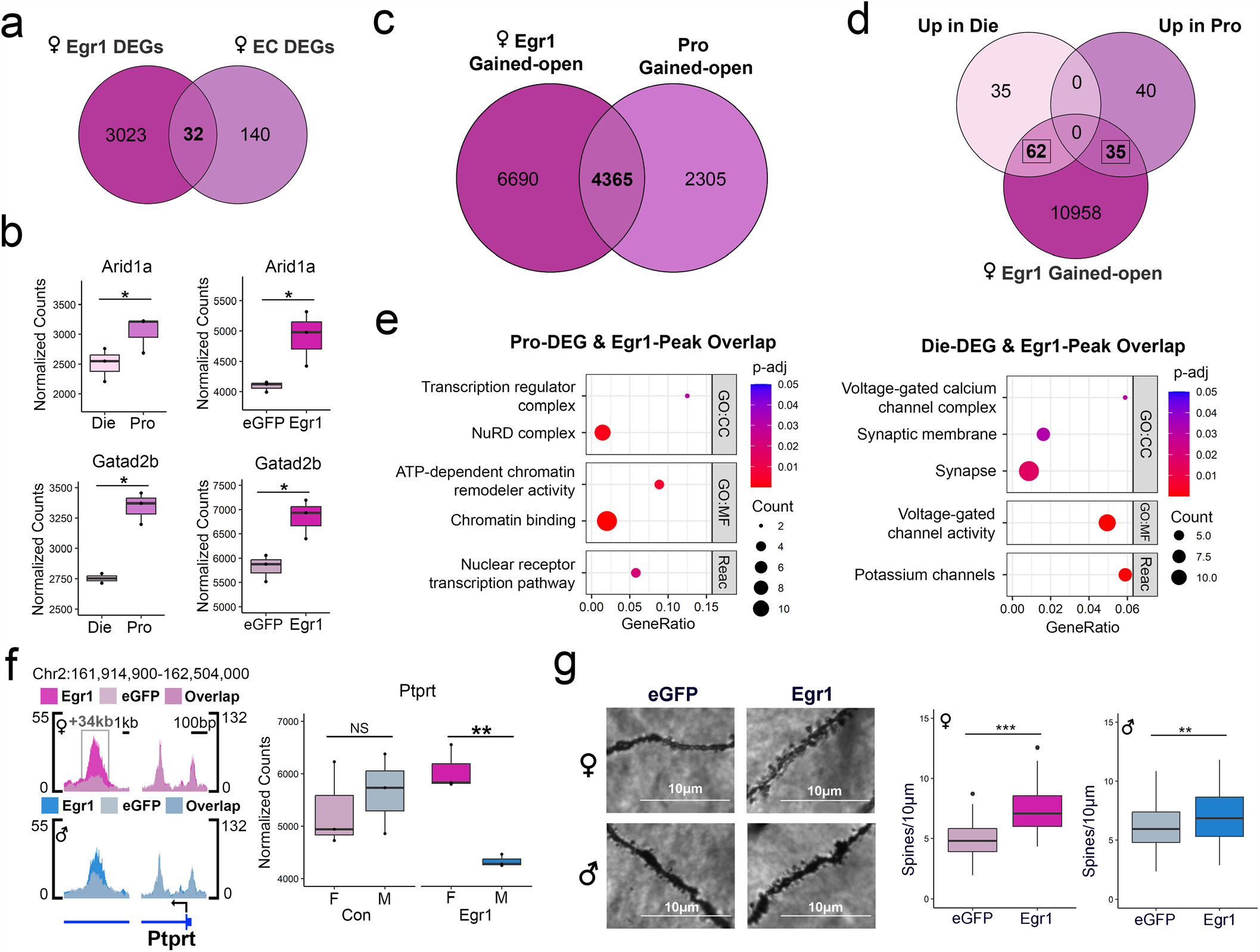
Egr1 overexpression partially recapitulates proestrus-associated changes in gene expression, chromatin, and synaptic plasticity. **a**. There is a significant overlap of Egr1 overexpression-induced and oestrous cycle (EC)-related DEGs. **b**. Genes upregulated in proestrus and induced by Egr1 overexpression include *Arid1a* and *Gatad2b*. **c**. There is a significant overlap of female Egr1-gained open chromatin regions and proestrus (Pro)-gained open chromatin regions. **d**. Oestrous cycle-related DEGs exhibit an even higher overlap with genes bearing Egr1 gained-open regions. **e**. The overlapping genes that are induced in proestrus and those that are downregulated in proestrus are enriched for different Gene Ontology (GO) terms. Dot plots colours indicate p-values and dot size corresponds to the number of overlapping genes between the list and the term/pathway (Count); the x-axis corresponds to the ratio of genes in the input list to genes in the term/pathway (GeneRatio). **f**. *Ptprt* is a synaptic function-related gene that shows sex-specific changes in chromatin accessibility and gene expression following Egr1 overexpression. ATAC-seq data is shown using SparK plots of group-average normalized ATAC-seq reads (n = 4 biological replicates/group/sex). RNA-seq data is shown using count plots (n=3 biological replicates/group/sex). Box plots (box, 1st-3rd quartile; horizontal line, median; whiskers, 1.5xIQR); RNA-seq significance codes; ^*^, Padj<0.05; ^**^, Padj<0.01; NS, non-significant; Benjamini-Hochberg correction for multiple testing. **g**. Egr1 overexpression increases dendritic spine density in the CA1 vHIP neurons of both OVX females and males, but the effect size is higher in females (d=1.532) compared to males (d=0.4468). Dendritic spine density significance codes; ***, P<0.001; **, P<0.01; Welch two-sample t-test. Die, dioestrus; Pro, proestrus; CC, Cellular Component; MF, Molecular Function; Reac, Reactome; F, female; M, male.

### Egr1 overexpression partially recapitulates proestrus-associated chromatin changes in vHIP neurons

We next performed overlaps of chromatin accessibility between the oestrous cycle^9^ and Egr1 overexpression experiments. Interestingly, compared to gene expression, we see a much greater overlap between the ATAC-seq datasets, where 77% (7402/9614) of genes with differential chromatin peaks across the oestrous cycle also show changes in chromatin in OVX females after Egr1 overexpression (**Extended Data Fig. 8**). Since we specifically implicated Egr1 in mediating the proestrus behavioural and transcriptional state^9^, we next overlapped Egr1 gained-open regions with proestrus gained-open regions and found 65.4% (4365/6670) of genes with proestrus gained-open chromatin also exhibit gained-open chromatin after Egr1 overexpression (**Figure 4c**). Due to the extensive effects of Egr1 on chromatin accessibility and the larger overlaps between the Egr1 overexpression and oestrous cycle ATAC-seq datasets, we explored the overlap between Egr1’s chromatin effects and oestrus cycle DEGs. The reasoning behind this was that Egr1’s directed effects on chromatin may be more representative of its physiological effects across the cycle than its effects on gene expression, which likely rely on the presence of additional transcription factors that may be absent in the vHIP neurons of females that underwent ovariectomy. This analysis revealed that 56.4% (97/172) of oestrous cycle DEGs have Egr1 gained-open chromatin after Egr1 overexpression in OVX females (**Figure 4d**). Of these DEGs, 64.9% (63/97) also have chromatin changes across the oestrous cycle, indicating the importance of chromatin accessibility in their physiological regulation (**Extended Data Fig. 9**). Consistent with Egr1’s role in both activating^34^ and repressing transcription^35^, we found Egr1 gained-open regions overlap with genes both up- and downregulated in proestrus compared to dioestrus females (**Figure 4d**). Overlapping genes upregulated in proestrus (down in dioestrus) are enriched for terms and pathways related to chromatin and transcription regulation, while overlapping genes downregulated in proestrus (up in dioestrus) are enriched for terms related to synaptic function and ion channels (**Figure 4e**). These results indicate that Egr1 mediates opening of chromatin near its binding motifs near genes that play a critical role in transcriptional regulation and neuronal function within physiologically cycling female mice.

### Egr1 overexpression recapitulates proestrus-associated changes in synaptic plasticity in vHIP neurons

Notably, we found enrichment of genes relevant to synaptic function among female-specific Egr1-induced DEGs (**Figure 2e**) and in the overlap between Egr1’s chromatin effects and oestrous cycle DEGs **(Figure 4e**). Thus, we hypothesized that Egr1’s gene regulatory effects may drive sex-specific changes in the density of dendritic spines, the sites of active synapses. As an example, *Ptprt* (encoding Protein tyrosine phosphatase receptor T) is a putative Egr1 target gene that is implicated in the regulation of dendritic spines^36^, and was previously shown to exhibit increased expression and a more open chromatin state in vHIP neurons during the proestrus (compared to dioestrus) phase of the oestrous cycle^9^. Notably, Egr1 overexpression mimics proestrus, leading to an increase in chromatin accessibility at an intronic region of *Ptprt* in females (but not in males; **Figure 4f**). Further, Egr1 induction induces a sex difference in *Ptprt* expression, with females exhibiting higher expression than males following Egr1 overexpression which we did not observe in controls (**Figure 4f**).

Following up on this sex-specific regulation of genes involved in synapse regulation, we next assessed whether Egr1 overexpression alters dendritic spine density in vHIP neurons. After delivering Egr1 or control AAVs to males and OVX females, we performed Golgi-Cox staining and imaged dendrites of ventral CA1 pyramidal neurons. We found that Egr1 overexpression increases vHIP dendritic spine density in OVX females (**Figure 4g**), again mimicking the proestrus phase of the cycle^9^. Importantly, Egr1 overexpression also increased the spine density in males, but the effect size was considerably larger in females (d = 1.532) than in males (d = 0.4468).

## Discussion

In summary, our study reveals that Egr1 is a sex-specific IEG, capable of opening neuronal chromatin sex-specifically, with consequences for targeted gene expression changes, synaptic plasticity, and anxiety- and depression-related behaviour. This role of Egr1 in directing chromatin accessibility is consistent with its previously established epigenetic regulatory role in recruiting Tet1 to shape the brain methylome in response to neuronal activity^37^. In addition, this is reminiscent of the role of cFos, another IEG, shown to direct chromatin opening in a neuronal activity-dependent manner^17^. However, the notion of sex-specific IEGs is novel to neuroscience, and our data demonstrate that IEGs such as Egr1 allow for sex hormone-driven chromatin and brain plasticity. This response to hormones seems to be optimized by oestrogen’s regulation of Egr1 expression, as reflected in the cyclical Egr1 expression pattern across the oestrous cycle in the vHIP, but also in other tissues^38,39^. In fact, this cyclical expression is indicative of Egr1’s role in driving the biological clocks in the body, since Egr1 is also induced in the suprachiasmatic nucleus by light stimulation^40^, and has been implicated in the circadian rhythm-driven plasticity^19,41^. Finally, neuronal Egr1 expression has been shown to vary with stress^42,43^, psychiatric disease^44,45^, and antidepressant treatment^46^, and therefore represents a master molecular adaptor at the interface of the external and internal environment, with a critical role in sex-specific neurobiology and disease risk.

Importantly, we were able to go from genomics data generated using the physiological mouse model as a function of the oestrous cycle to *in vivo* mouse manipulation studies to validate a sex-specific regulator of chromatin, neuroplasticity and behaviour. Since ovarian hormone shifts are an established but underexplored psychiatric risk factor in humans^6,7,47-50^, our study provides a path to find new sex-based drug targets and treatments that could be transformational for neuropsychiatric disorders and women’s brain health in particular.

## Acknowledgements

This work was supported by the National Institutes of Health award R01MH123523 (to M.K.). We would further like to acknowledge the resources of the Center for Epigenomics at the Albert Einstein College of Medicine. Finally, we would like to thank Ming Liu and Lara Winterkorn for their assistance with nuclei sorting and next generation sequencing, respectively. Schematics in Figures 1, 2, and 3, and in Extended Data Figures 1 and 6 were created using BioRender.com.This paper was typeset with the bioRxiv word template by @Chrelli: www.github.com/chrelli/bioRxiv-word-template

## Author contributions

D.R. and M.K. designed the study; D.R., E.P., E.F.G., and M.K. performed experiments. D.R. and M.S. performed data analyses. D.R. and M.K. interpreted the data. D.R. constructed the figures. J.M.G. contributed computational resources. E.F.G. contributed to AAV vector design. D.R. and M.K. wrote the article. M.K. conceived and directed the project. All authors commented on and approved the final version of the paper.

## Data and materials availability

RNA-seq and ATAC-seq data will be available on GEO upon publication of the manuscript. All other materials are included in the manuscript or are available from the authors upon request. Correspondence and requests for materials should be addressed to M.K.

## Materials and Methods

### Animals

For this study we used five separate cohorts of animals. The first cohort was used for behavioural and gene expression analysis of intact females across the oestrus cycle and was comprised of n=48 C57BL/6J female mice that arrived at 5 weeks of age from Jackson Laboratory. The second cohort was used to compare the behaviour of gonadally intact and ovariectomized female mice and included n=34 intact and n=12 ovariectomized C57BL/6J female mice, with ovariectomies performed at the Jackson Laboratory at 4 weeks of age. Ovariectomized mice were allowed to recover for one week before arriving, together with the intact females, at 5 weeks of age. After habituating for two weeks (5-7 weeks old), intact females from these cohorts underwent oestrous cycle tracking daily in the morning (between 9AM and 11AM) for two weeks (7-9 weeks) in order to establish a predictive cycling pattern for each female animal and to ensure that only females with regular cycles were included in the study (see *Oestrous cycle monitoring*). Vaginal smears were also taken from ovariectomized females for 4 days to verify that ovariectomy was successful. This was followed by behavioural testing at 9-11 weeks of age (see *Behavioural testing*). Animals were sacrificed at 11 weeks of age by cervical dislocation, brains were extracted, and bilateral ventral hippocampi were dissected on ice then flash frozen in liquid nitrogen.

The remaining three cohorts were used in the overexpression experiments and included ovariectomized C57BL/6J females alongside age-matched males. These cohorts were used for behaviour (n=24/sex), molecular analyses (n=20/sex), and Golgi staining (n=10/sex), respectively. Females underwent ovariectomy at the Jackson Laboratory at 4 weeks of age and recovered for 1 week before arriving at 5 weeks of age. Males similarly arrived at 5 weeks of age. All animals were allowed to habituate before undergoing stereotaxic surgery (see *Stereotaxic surgery*) at 7 weeks of age and were allowed to recover ∼3 weeks to ensure stable transgene expression. During recovery, 4 days of vaginal smears were obtained from ovariectomized females to confirm ovariectomy. In the behaviour cohort, animals underwent behavioural testing from 10-11 weeks of age before being sacrificed at 11 weeks of age, at which time the whole brain was removed and preserved for cryosectioning (see *Immunohistochemistry*) so that we could verify viral targeting in every animal that underwent behavioural testing. For the molecular analysis cohort, animals were sacrificed at 10 weeks and bilateral hippocampi were dissected on ice and flash frozen in liquid nitrogen. Analysis of RNA-seq data in this cohort confirmed overexpression of Egr1. For the Golgi staining cohort, animals were sacrificed at 10 weeks and the whole brain was removed then underwent Golgi-Cox staining (see *Golgi-cox staining for analysis of dendritic spines*).

All animals used in this study were housed in same-sex cages (n=3-5/cage) and were kept on a 12:12h light:dark cycle with *ad libitum* access to food and water. All frozen tissues were stored at -80°C before further processing. All animal procedures were approved by the Institutional Animal Care and Use Committee at Fordham University.

### Oestrous cycle monitoring

Oestrous cycle tracking was performed using vaginal smear cytology, a well-established method, as described previously^1-3^. Smears were collected by filling a disposable transfer pipette with 100 µl of distilled water, gently placing the tip of the pipette at the vaginal opening and collecting cells via lavage. The cell-containing water was then applied to a microscope slide and allowed to dry at room temperature for two hours. Once dried, slides were stained with 0.1% crystal violet in distilled water, washed, and then allowed to dry prior to examination with light microscopy. Oestrous cycle stage can be determined by the relative quantities of nucleated epithelial cells, cornified epithelial cells, and leukocytes. The proestrus phase is characterized by mostly nucleated epithelial cells while the oestrus phase is characterized by mostly cornified epithelial cells. Metestrus and dioestrus have both forms of epithelial cells as well as leukocytes, and dioestrus exhibits a higher proportion of leukocytes compared to metestrus. After determining the oestrous cycle stage of each female animal daily for two weeks, or three cycles, cycling patterns can be established and stage predictions can be made for the grouping of animals for behavioural tests and molecular analyses. Grouping predictions were confirmed by obtaining vaginal smears of animals following behavioural testing or sacrificing. We previously validated that our predictions correspond to the expected ovarian hormone content in both serum and hippocampal tissue for proestrus (high-oestrogen, low-progesterone) and dioestrus (low-oestrogen, high-progesterone) mice^2^. Animals with irregular cycles were excluded from behavioural and molecular analyses.

### Candidate gene expression

To analyse the gene expression of *Egr1* in cycling, ovary-intact animals, RNA was isolated from the ventral hippocampus with Qiagen’s Allprep DNA/RNA Mini Kit (n=24 females, 6 per oestrous cycle stage). Reverse transcription was then performed with Invitrogen’s SuperScript III First-Strand Synthesis kit, and qPCR was performed with Applied Biosystems’ FAST SYBR Green and the Quant Studio 3 PCR machine. Relative *Egr1* and *Ppp1r1b* transcript levels were determined using the 2-ΔΔCt method with oestrus as the reference group and Cyclophilin A (*Ppia*) as the endogenous reference gene. The following primer sequences were used in this analysis: *Egr1* (forward: 5′-AGCGAACAACCCTATGAGCAC-3′, reverse: 5′-GGA-TAACTCGTCTCCACCATCG-3′); *Ppp1r1b* (forward: 5′-GGACGAAGAAGAAGA-CAGCCA-3′, reverse: 5′-CACTTGGTCCTCAGAGTTTCC-3′); *Ppia* (forward: 5′-GAGCTGTTTGCAGACAAAGTTC-3′, reverse: 5′-CCCTGGCACATGAATCCTGG-3′).

### Behavioural testing

Animals underwent behavioural testing from 9-11 weeks of age. For all behavioural tests, the movement of animals was tracked with a camera and AnyMaze© tracking software. For the open field test, animals were placed in the corner of a 40 x 40 x 35cm open field arena (Stoelting Co.). The animal’s movement was tracked for 10 minutes and the total distance travelled by the animal, the time spent in the centre versus the periphery of the maze, and the number of entries into the centre were recorded. For the elevated plus maze test, animals began the test by being placed in the centre of the raised (40cm) plus-shaped platform (two 35 x 5cm arms; Stoelting Co.). The movement of the animal was tracked for 5 minutes and the total distance travelled by the animal as well as the time spent in the open arms versus the closed arms was recorded. For the forced swim test, animals were placed in a 2 L glass beaker with 1 L of lukewarm water. Movement was tracked for 2 minutes and the time spent immobile was recorded.

### Adeno-associated viruses

Recombinant adeno-associated viruses (AAVs) were used to overexpress Egr1 in ventral hippocampal neurons. The AAV9 serotype was selected due to its suitability for gene transduction in brain tissue^4^.The two AAVs used in these experiments were an experimental AAV and a control AAV; the experimental AAV contains the Egr1 transgene, an IRES element, and eGFP, with the IRES element ensuring coproduction of Egr1 and eGFP from a single AAV, while the control AAV contains only eGFP. Both control and experimental AAVs utilize the neuron-specific hSYN promoter^5^. Both AAVs also contain a WPRE enhancer used to increase the expression of the respective transgenes, with the experimental AAV containing a smaller iteration of this enhancer (WPRE3) due to space limitations which retains a high degree of enhancer activity^6^.Both viruses were purchased from Vector Biolabs and were provided in PBS solution with 5% glycerol at a titre of ∼10^13^ genome copies/ml.

### Stereotaxic surgery

Animals were anaesthetized prior to surgery using isoflurane delivered to an anaesthesia induction chamber and kept anaesthetized during surgery with a continuous flow of 1.5-2% isoflurane. Either control or experimental AAV solution (n=12/group/sex for behaviour, n=6/group/sex for RNA-seq, n=4/group/sex for ATAC-seq, n=5/group/sex for Golgi staining) was delivered to the injection site through a glass pipette (15μm diameter) fixed to a Nanoject III© microinjector pump. The coordinates (relative to bregma) used for ventral hippocampal injections were: A/P: -2.95, M/L: +/-2.85 (bilateral injections), D/V: -3.8, -3.9. 200nL was delivered (10 pulses of 20nL with 15 seconds in between) at D/V -3.9, then again at D/V -3.8 totalling 400nL of injection/side. Three weeks were allowed to pass to permit stable expression levels of the viral transgenes before behavioural testing was conducted or before the animals were sacrificed for molecular analyses or Golgi staining. In order to verify proper viral delivery into the ventral hippocampus of the animals that underwent behavioural testing, immunostaining of brain sections was performed (see *Immunohistochemistry*). Animals were excluded if there was no viral expression present or if there was evidence of off-target expression or tissue damage (this occurred in n=4 females and n=2 males, or 12.5% of animals tested). For molecular analyses, we were able to confirm efficient Egr1 overexpression in ventral hippocampal neurons using Egr1 transcript levels (RNA-seq) and chromatin accessibility data (ATAC-seq).

### Immunohistochemistry

Animals were sacrificed by cervical dislocation and brains were removed. Brains were then washed with ice cold 0.1M PBS and fixed in 4% PFA in 0.1M PBS at 4°C for 24 h. After fixation, brains were rinsed in cold 0.1M PBS and underwent sucrose preservation, which involved placing the brains in solutions containing 15% then 30% sucrose dissolved in 0.1M PBS at 4°C for 24h and 48h, respectively. Brains were then frozen in dry ice-cooled hexane and stored at -80°C until sectioning. Cryosectioning was performed by embedding brains in optimal cutting temperature compound (OCT) and cutting serial sections on a rotary cryostat (Leica CM1950, Leica Biosystems GmBH). 20 μm coronal sections containing the ventral hippocampus were collected on Super Frost Ultra Plus slides (Fisher Scientific) and stored at -80°C prior to immunostaining. Immunostaining involved rehydrating slides in 0.1M PBS for 30 minutes at room temperature. After rehydration, blocking buffer (5% BSA and 0.4% Triton X-100 in 0.1M PBS) was applied for 1 hour at room temperature. Slides were then washed with PBS-T (1% BSA and 0.4% Triton X-100 in 0.1M PBS), and the Egr1 primary antibody was applied (Rabbit anti-Egr1 mAb Cell Signal #4153, 1:500 in PBS-T) for 24h at 4°C. Following primary antibody incubation, slides were washed with PBS-T, then the secondary antibody was applied (Donkey anti-Rabbit IgG pAb Invitrogen A-21207, 1:250 in PBS-T) for 2 hours in the dark at room temperature. Following secondary antibody incubation, slides were washed with PBS-T, then DAPI was applied (1:1000 in PBS) for 5 minutes in the dark at room temperature. Following DAPI incubation, slides were washed with 0.1M PBS and mounted with Mowiol 4-88 and a coverslip. For all staining sessions, sections with no primary antibody (PBS-T applied instead of primary antibody) and sections with no secondary antibody (PBS-T applied instead of secondary antibody) were included to ensure that the observed fluorescent signal corresponded to Egr1 detection rather than autofluorescence of the tissue or non-specific binding of the secondary antibody.

### Fluorescence-activated nuclei sorting (FANS)

For overexpression experiments, all molecular analyses were performed on neuronal nuclei which were purified using FANS, as described previously^7,8^. Briefly, for the ATAC-seq experiment, 4 animals per group (eGFP, Egr1) per sex were included (total n = 16). For the nucRNA-seq experiment, 6 animals per group (eGFP, Egr1) per sex were included (total n = 24). The ATAC-seq experiment used bilateral ventral hippocampi from a single animal for each biological replicate (n=4/group/sex), while for nucRNA-seq, bilateral ventral hippocampi were pooled from two animals for each biological replicate (n = 3 replicates/group/sex). We performed FANS in batches of 3-4 biological replicates per session and ensured that the groups were evenly dispersed across batches for each experiment. Brain tissue was dissociated in lysis buffer using a tissue douncer and nuclei were isolated from tissue lysates by ultracentrifugation through a sucrose gradient. Pelleted nuclei were resuspended in DPBS and incubated for 45 minutes with the mouse monoclonal antibody against neuronal nuclear marker NeuN conjugated to AlexaFluor 488 (1:1000, MAB377X; Millipore, MA). Before sorting, we added DAPI (1:1000) to the incubation mixture and filtered all samples through a 35-µm cell strainer. FANS was performed on a FACSAria instrument (BD Sciences) and data were collected and analysed using BD FACSDiva v8.0.1 software at the Albert Einstein College of Medicine Flow Cytometry Core Facility (**Extended Data Fig. 10**). In addition to a sample containing NeuN-AlexaFluor 488 and DAPI stain, three controls were used to set up the gates for sorting: DAPI only; IgG1 isotype control-AlexaFluor 488 (1:1000, Mouse monoclonal IgG1-k, FCMAB310A4; Millipore, MA) and DAPI; and NeuN-AlexaFluor 488 only. We set up the protocol to remove debris, ensure single nuclear sorting (using DAPI), and select the NeuN+ (neuronal) and NeuN-(non-neuronal) nuclei populations (Extended Data Fig. 10). For the ATAC-seq experiment, we collected 50,000 NeuN+ nuclei per biological replicate in BSA-precoated tubes filled with 200 µL of DPBS. For the nucRNA-seq experiment, we collected 142,041-250,000 nuclei directly into Trizol LS reagents (Thermo Fisher Scientific, 10296-010) to protect RNA from degradation.

### nucRNA-seq

Following sorting of nuclei, chloroform was added to the sample and the aqueous phase was recovered. nucRNA was then isolated and purified using the RNeasy Micro kit (Qiagen). The quality of nucRNA was assessed using the Fragment Analyzer (Agilent), with each sample having RQN>7 (**Extended Data Fig. 11a**). nucRNA quantity was determined using the Qubit RNA High-Sensitivity assay (Thermo Fisher Scientific, Q32852). All purified nucRNA (9-20 ng/sample) was used to construct cDNA sequencing libraries using the KAPA RNA HyperPrep Kit with RiboErase (KK8560, KAPA Biosystems), following the manufacturer’s instructions. rRNA was depleted from nucRNA samples using hybridization of DNA oligonucleotides complementary to rRNA followed by RNase H and DNase treatment. The efficiency of rRNA depletion was confirmed using the Bioanalyzer (Agilent) and with qRT-PCR amplification of the 28S rRNA transcript before and after RiboErase treatment using the following primers: Forward, 5’-CCCATATCCGCAGCAGGTC-3’; Reverse, 5’-CCAGCCCTTA-GAGCCAATCC-3’. rRNA-depleted nucRNA was then fragmented at 94°C for 5 minutes in the presence of Mg2+, followed by first and second strand cDNA synthesis and A-tailing. After ligation of KAPA dual-indexed adapters (KK8722), each library was amplified using 13-14 PCR cycles. Following a bead-based clean-up, the length of the libraries (365bp, on average) and their quality was assessed using the Bioanalyzer High-Sensitivity DNA Assay (**Extended Data Fig. 11b**). The libraries were quantified with the Qubit HS DNA kit (Life Technologies, Q32851) and by qPCR (KAPA Biosystems, KK4873) prior to sequencing. 150 bp, paired-end sequencing was performed in one lane of an S1 flow cell on the NovaSeq 6000 instrument at the New York Genome Center, yielding 60-90 million reads per sample.

### RNA-seq analysis

Sequences were adapter-trimmed and aligned to the mouse reference genome (mm39) with the GENCODE M32 gtf file using Star Aligner^9^. The gene counts produced by Star were collated into a matrix. Differential gene expression analysis was then performed using DESeq2^10^ at significance level p_adj_<0.05. Volcano plots were created using EnhancedVolcano (https://github.com/kevinblighe/EnhancedVolcano). Enrichment analyses were performed using FUMA^11^ at significance level p_adj_<0.05. Enrichment analysis was also performed using GSEA^12^ with ranked gene lists and results were plotted using the EnrichmentMap^13^ and AutoAnnotate apps in Cytoscape^14^.

### ATAC-seq

We performed ATAC-seq according to Buenrostro et al.^15^ and our previously published protocol on sorted neuronal nuclei^2,8^, with some modifications^17^. Following FANS, nuclei were pelleted and resuspended in 50 µL of the transposase reaction mix including 25 µL of the 2xTD reaction buffer, 5 µL of transposase enzyme (Illumina Tagment DNA TDE1 kit, 20034197). We also included additional components: 16.5µl 0.1M PBS, 0.5µl 1% digitonin, and 0.5 µl 10% Tween-20, which were previously shown to enhance transposition efficiency^17^. Transposition occurred at 37°C for 30 minutes and transposed DNA was purified using the MinElute PCR Purification Kit (Qiagen, 28004). Indexing libraries using Nextera i5 and i7 indexed amplification primers (Illumina XT index kit, 15055290) occurred alongside library amplification with NEBNext High-Fidelity PCR Master Mix (New England Biolabs, M0541S). The PCR reaction was carried out with the following conditions: 1 cycle of 72°C for 5 minutes and 98°C for 30 seconds; followed by 10 cycles of 98°C for 10 seconds, 63°C for 30 seconds, and 72°C for 1 minute. Amplified libraries were then purified using the MinElute PCR Purification Kit. Library quality was then assessed using the Bioanalyzer High-Sensitivity DNA Assay (**Extended Data Fig. 11c**). The ATAC-seq libraries were quantified by Qubit HS DNA kit and by qPCR prior to sequencing. 150 bp, paired-end sequencing was performed in one lane of an S1 flow cell on the NovaSeq 6000 instrument at the New York Genome Center, yielding 40-75 million reads per sample.

### ATAC-seq analysis

Sequences were adapter trimmed and aligned to the mouse reference genome (mm39) using BWA-MEM software^18^. Bam files were down-sampled prior to peak-calling to ensure a comparable number of reads for each sample. The peak-calling was performed using MACS2 as previously reported^15^. We ensured all samples had greater than 20% of reads in peaks, indicative of high-quality ATAC-seq samples^8^. High confidence peaks were then selected for downstream analysis by overlapping the replicates in each group (n = 4/group/sex) and continuing the analysis with peaks that were called in at least 3/4 replicates. Peak annotation was performed using the annotatePeak-InBatch function on UCSC.mm39.refGene annotation^19^, and we identified overlapping peaks using the findOverlapsOfPeaks function of ChIPpeakAnno^20^. Motif analysis was performed using HOMER^21^ and enrichment analysis on lists of annotated genes was performed with EnrichR^22^. Plots of aligned ATAC-seq data were created using SparK^23^. Heatmaps and profile plots of H3K4me1, H3K27ac, and H3K4me3 enrichment within gained-open Egr1 binding sites was performed using deepTools^24^ using the publicly-available ChIP-seq data <u>GSE74964</u>^25^ generated using neuronal (NeuN+) CA1 hippocampal neurons from male mice.

### Integration of genomics data

Our ATAC-seq and nucRNA-seq data previously generated from ventral hippocampal neurons <u>(GSE114036</u>) of proestrus female, dioestrus female, and male mice^2^ were integrated with the data used in this study. For the purposes of overlapping the RNA-seq data, we applied the significance threshold used in the previous study^2^ (p_nominal_ < 0.05). Plots of aligned ATAC-seq data were created using SparK^23^.

### Golgi-cox staining for analysis of dendritic spines

Whole brains of mice injected with either control or Egr1 AAV (see *Adeno-associated viruses*, n=5/sex/group) were rinsed with distilled water and underwent the Golgi-Cox staining procedure (Golgi-Cox OptimStain Kit, Hitobiotec Inc. #HTKNS1125) following the manufacturer’s instructions. Briefly, whole brains were submerged in premixed Golgi-Cox impregnation solution and stored in the dark at room temperature for 24hrs, after which the solution was replaced and the brains remained submerged for an additional two weeks. Samples were then transferred to a tissue-protectant solution and kept in the dark at 4°C for 12 hours, after which the solution was changed and the brains remained submerged for an additional 72 hours. Samples were then frozen in dry ice-cooled hexane and stored at -80°C until sectioning. 100μm sections of OCT-embedded brains were cut using a cryostat (Leica CM1950) and collected on gelatin-coated slides. After drying overnight, slides were stained according to the kit protocol and mounted using DPX (Sigma-Aldrich). Brightfield Z-stacks of dendrites from ventral hippocampal CA1 pyramidal neurons were taken at 60x magnification and analysed using ImageJ (https://imagej.nih.gov/ij/index.html).

### Statistics

Statistics were performed using SPSS and R. For oestrous cycle gene expression and behavioural analysis, one-way ANOVA was used with Holm’s post hoc test. For behavioural and dendritic spine density analysis of animals from the overexpression experiments, Welch two-sample t-tests were performed. For each of these statistical tests, p < 0.05 was considered significant.

---

**Extended Data Figure 1.**
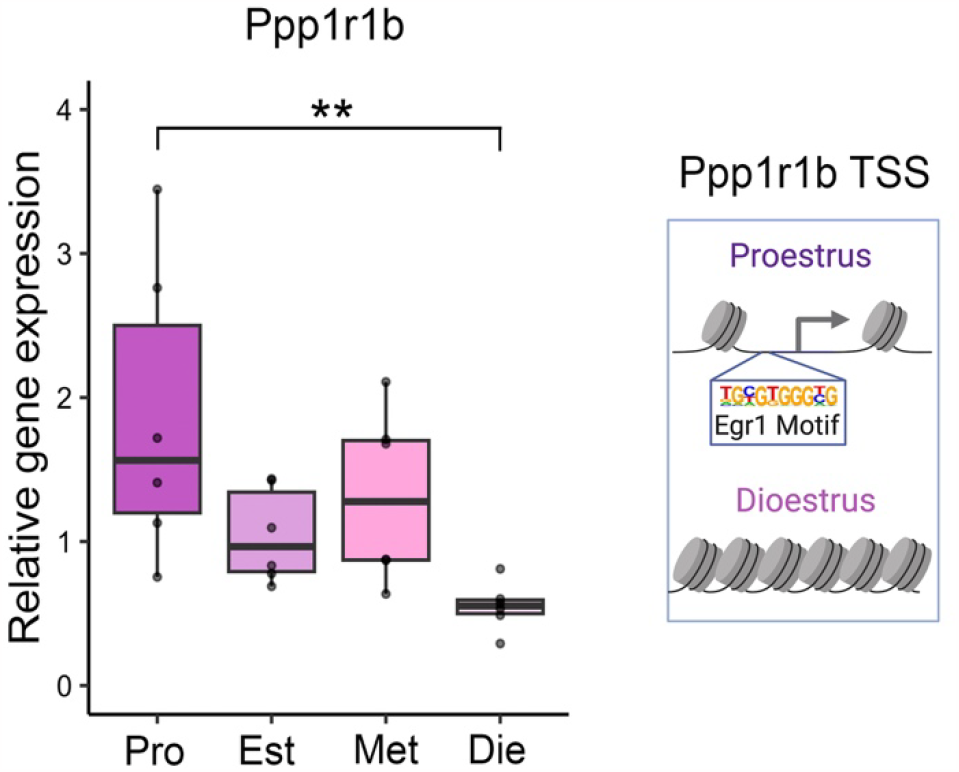
*Ppp1r1b* expression in the ventral hippocampus across the oestrus cycle. The expression of Egr1’s putative target gene *Ppp1r1b* changes in the ventral hippocampus over the oestrous cycle (left, n=5-6/group); the chromatin around the *Ppp1r1b* transcription start site (TSS), which contains the Egr1 binding motif, was previously found to be accessible in ventral hippocampal neurons of proestrus, but not dioestrus, female mice^9^ (right). Box plot (box, 1st-3rd quartile; horizontal line, median; whiskers, 1.5xIQR); one-way ANOVA with Holm’s post hoc test; **, P<0.01. Pro, proestrus (purple); Est, oestrus (light-purple); Met, metoestrus (pink); Die, dioestrus (light-pink).

**Extended Data Figure 2.**
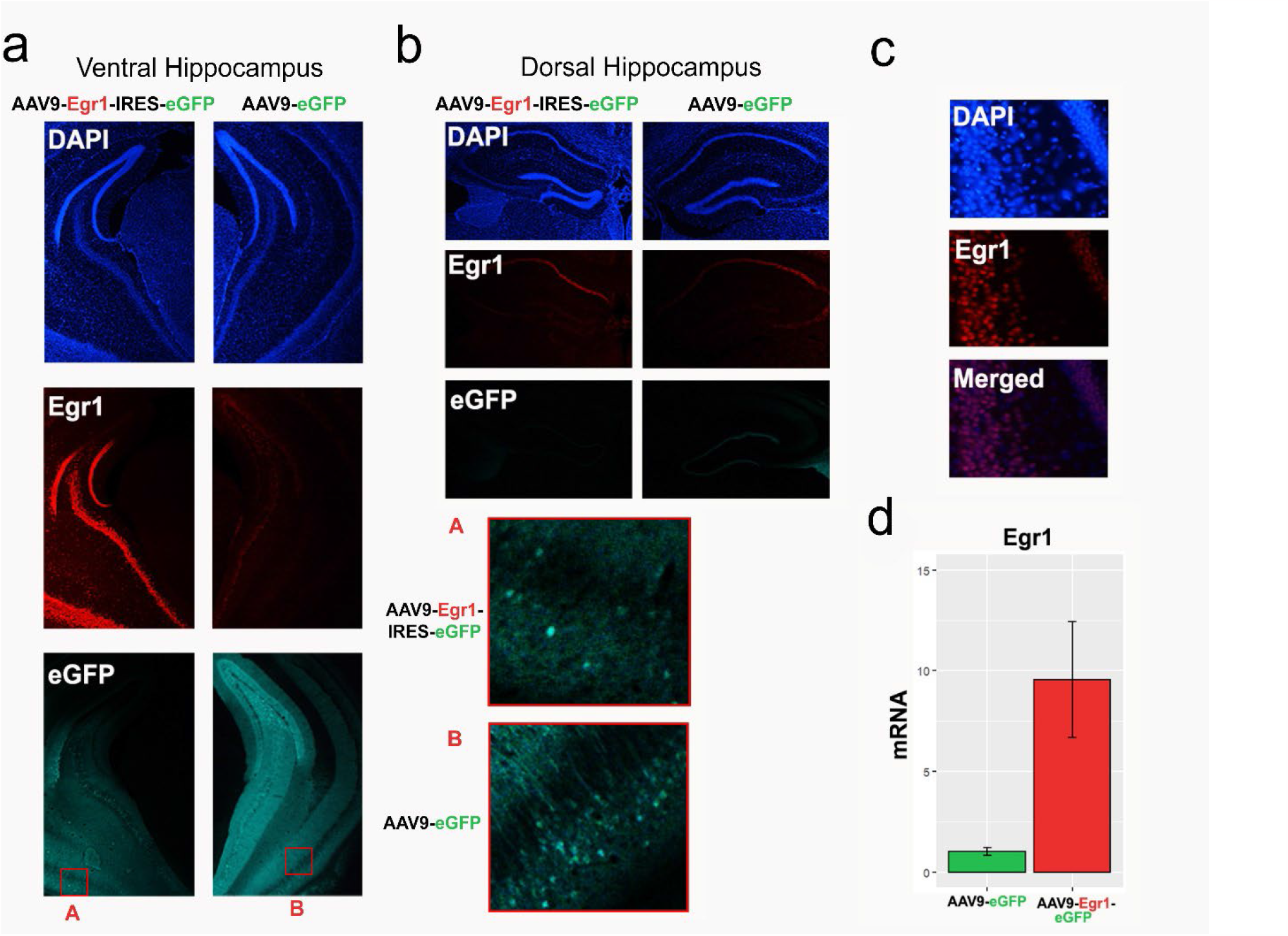
Verification of targeted Egr1 overexpression in the ventral hippocampus. **a-b**. Representative coronal sections containing the ventral (**a**) and dorsal (**b**) hippocampus were immunostained for Egr1 in animals that received either experimental (AAV9-Egr1-IRES-eGFP, left) or control (AAV9-eGFP, right) virus. Panels show DAPI (top), Egr1 (middle) and eGFP (bottom), with insets A and B (in red) demonstrating cellular eGFP expression in both sections. As targeted, we confirmed that Egr1 overexpression was observed in the ventral hippocampus (**a**) only, but not in the functionally distinct dorsal hippocampus (**b**). **c**. DAPI (top) and Egr1 (middle) signals overlap (bottom) in the ventral hippocampus of mice that received the Egr1 virus, confirming the nuclear localization of Egr1. Proper targeting of viral injections was verified for each animal, through either histology (behaviour cohort) or genomics data analysis (nucRNA-seq and ATAC-seq cohorts; see *Methods*) **d**. A pilot qRT-PCR experiment demonstrated *Egr1* mRNA overexpression (close to 10-fold) in the ventral hippocampus of animals injected with Egr1 virus, compared to animals that received the control virus (n=2/group), which was later confirmed with neuronal-specific nucRNA-seq (**Extended Data Fig. 5a**). Bar plots; whiskers denote 1.5x IQR.

**Extended Data Figure 3.**
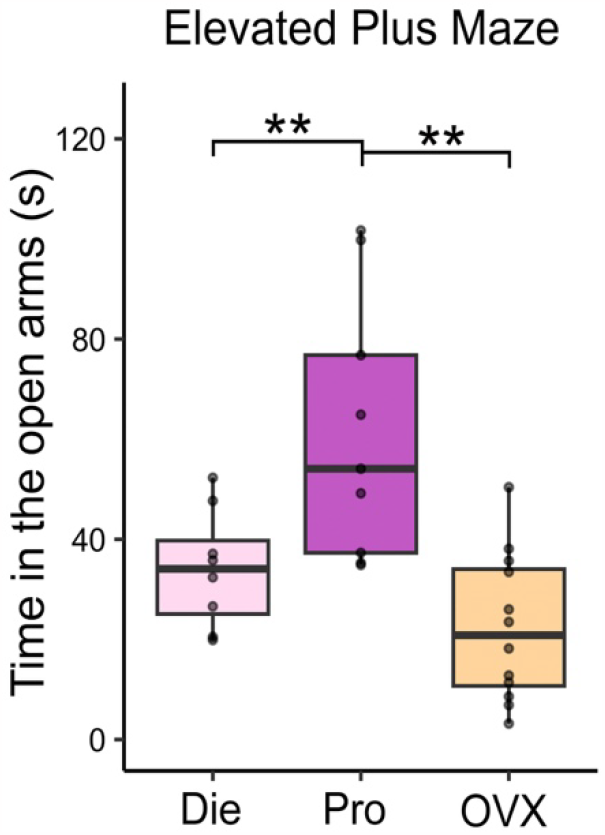
The effect of ovariectomy on anxiety-like behaviour. Ovariectomized (OVX) mice behave similarly to low-oestrogenic, dioestrus (Die) females and show higher anxiety indices than high-oestrogenic, proestrus (Pro) females, considering time spent in the open arms of the elevated plus maze (n=8-12/group). Box plot (box, 1st-3rd quartile; horizontal line, median; whiskers, 1.5xIQR); one-way ANOVA with Holm’s post hoc test; **, P<0.01. Die, dioestrus (pink); Pro, proestrus (purple); OVX, ovariectomized (yellow).

**Extended Data Figure 4.**
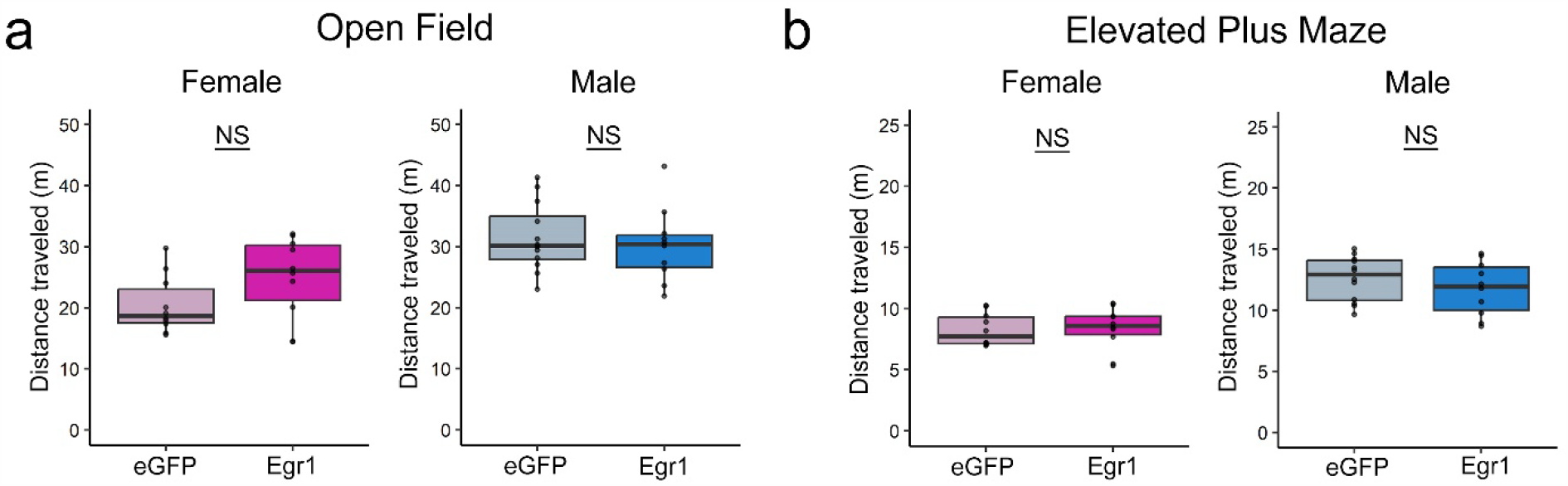
Total locomotor activity is not affected by Egr1 overexpression. Egr1 overexpression has no effect on distance travelled in either OVX females (left) or males (right) in the open field (**a**) or elevated plus maze (**b**) tests (n=10-12/group/sex). Box plots (box, 1st-3rd quartile; horizontal line, median; whiskers, 1.5xIQR); Welch two-sample t-test; NS, non-significant. Colour codes: eGFP females, pale pink; Egr1 females, bright pink; eGFP males, pale blue; Egr1 males, bright blue.

**Extended Data Figure 5.**
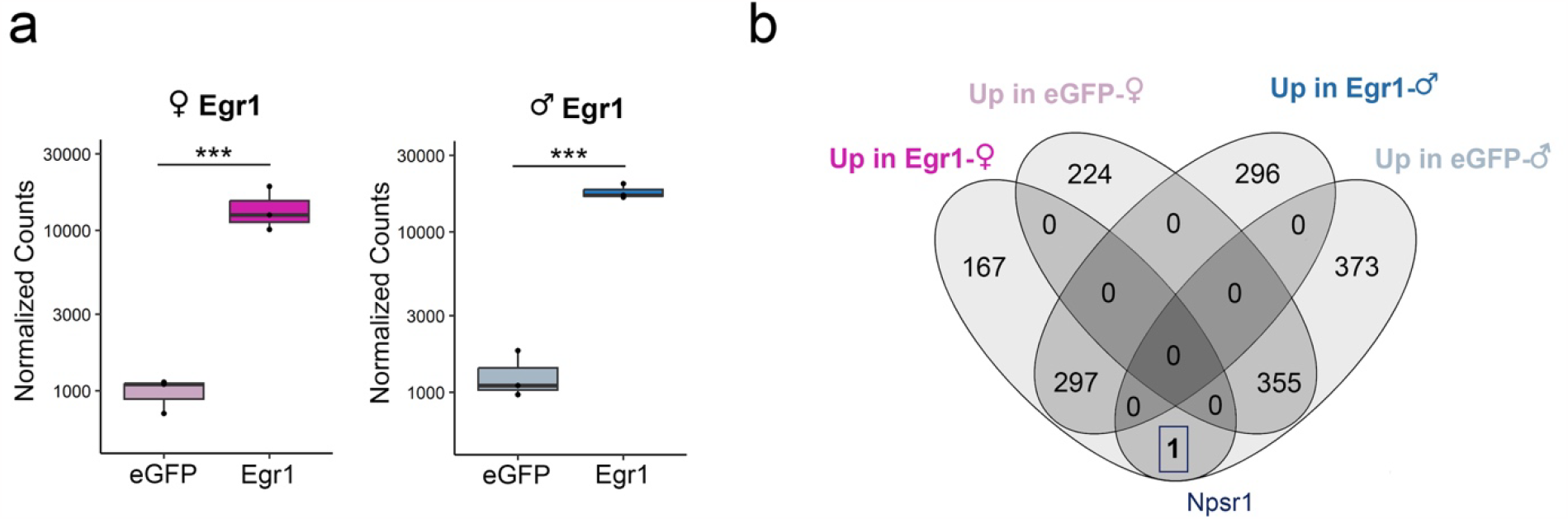
Egr1 overexpression and resulting sex-specific effects on gene expression in ventral hippocampal neurons. **a**. Count plots depicting transcript levels of *Egr1* in ventral hippocampal neurons of females (left) and males (right) following viral overexpression. **b**. A Venn diagram showing the overlap of differentially expressed genes between Egr1 and eGFP groups in females and males obtained by neuronal (NeuN+) nuclei specific RNA-seq analysis. The box denotes the *Npsr1* gene, which is downregulated in males and upregulated in females after Egr1 overexpression (**Figure 2f**). Box plots (box, 1st-3rd quartile; horizontal line, median; whiskers, 1.5xIQR). nucRNA-seq data show n=3 biological replicates/group/sex; ^***^, P_adj_<0.001; Benjamini-Hochberg correction for multiple testing. Colour codes: eGFP females, pale pink; Egr1 females, bright pink; eGFP males, pale blue; Egr1 males, bright blue.

**Extended Data Figure 6.**
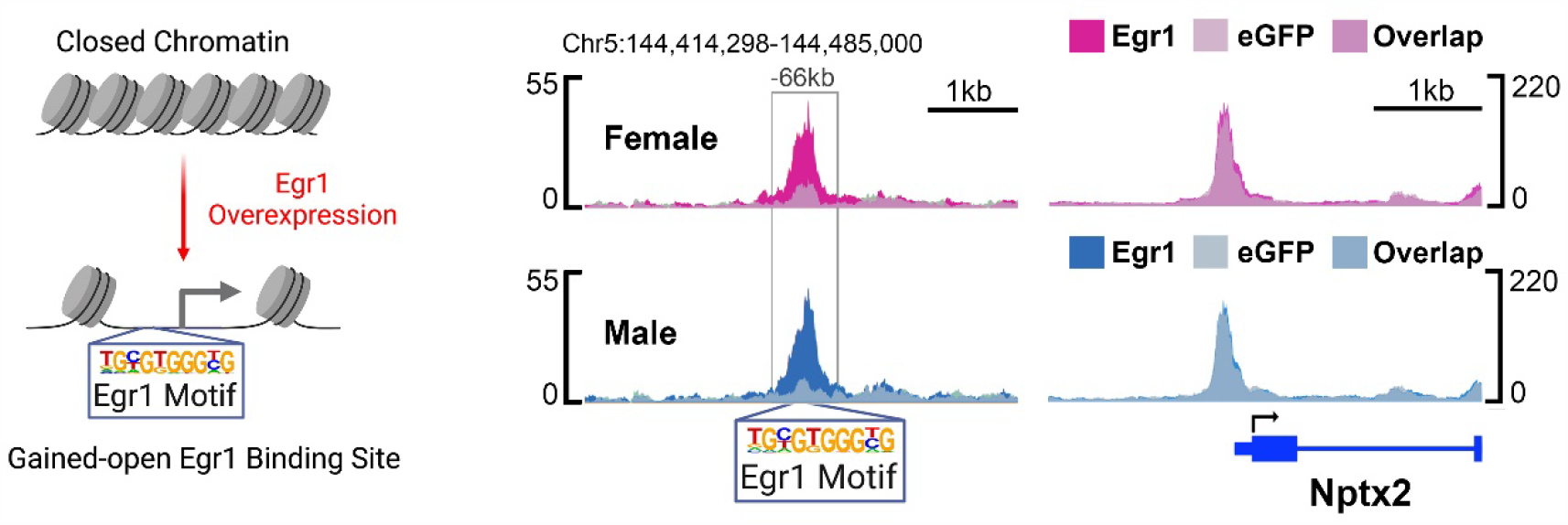
Egr1 gained-open chromatin in vHIP neurons. A scheme of Egr1-mediated opening of chromatin surrounding the Egr1 motif (left) in vHIP neurons and an example gene, *Nptx2* (right), which has an Egr1 gained-open putative enhancer region harbouring an Egr1 binding motif in both females (top) and males (bottom) 66kb upstream of the TSS. ATAC-seq data is shown using SparK plots of group-average normalized ATAC-seq reads (n = 4 biological replicates/group/sex). Colour codes: eGFP females, pale pink; Egr1 females, bright pink; eGFP males, pale blue; Egr1 males, bright blue.

**Extended Data Figure 7.**
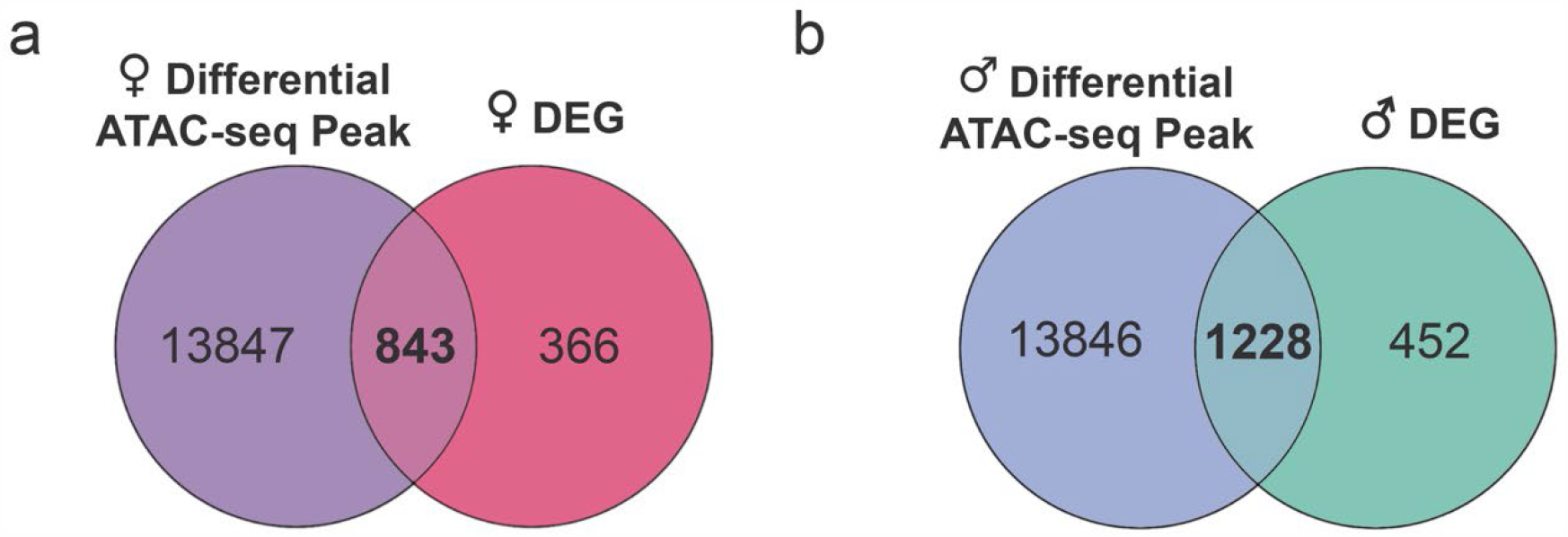
Egr1 has overlapping effects on chromatin accessibility and gene expression in ventral hippocampal neurons. **a-b**. A Venn diagram showing the overlap of genes annotated to Egr1-induced differential ATAC-seq peaks with Egr1-induced differentially expressed genes (DEGs) in ventral hippocampal neurons of females (**a**) and males (**b**).

**Extended Data Figure 8.**
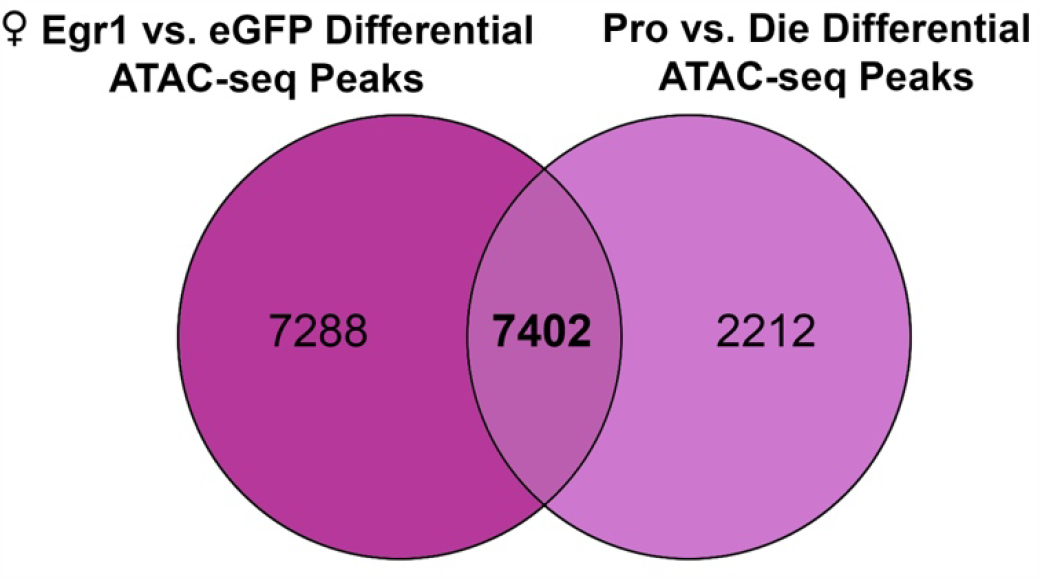
Egr1 induces overlapping chromatin organizational changes in ventral hippocampal neurons with proestrus females. A Venn diagram showing the overlap between genes annotated to female differential ATAC-seq peaks in the Egr1 overexpression and oestrous cycle experiments. Pro, proestrus; Die, dioestrus.

**Extended Data Figure 9.**
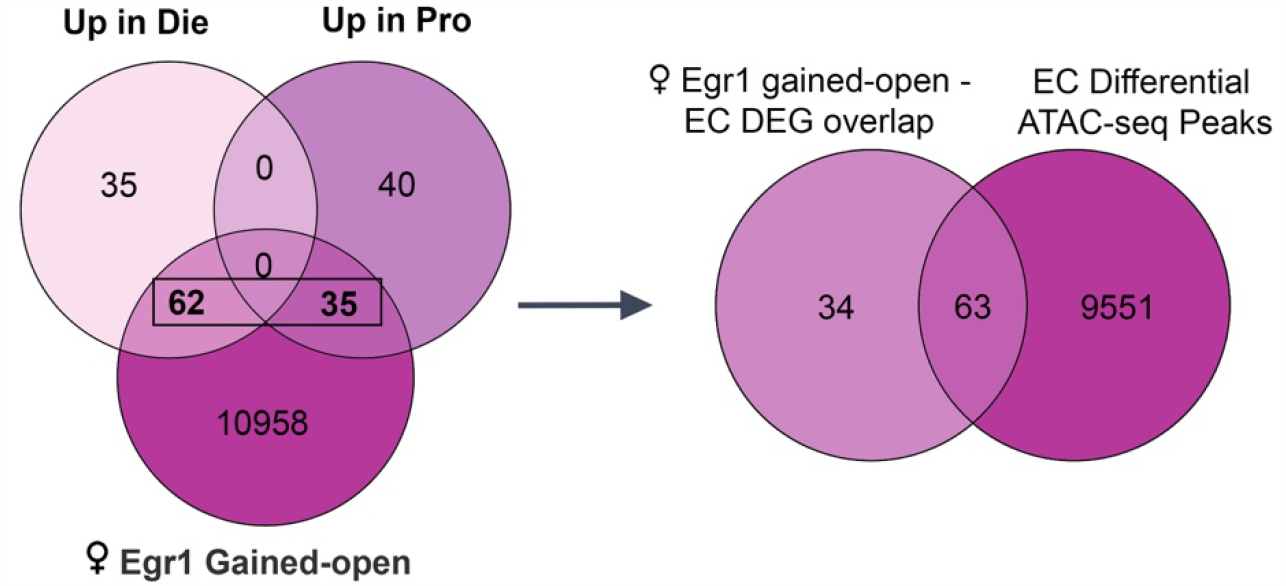
Overlapping elements of neuronal gene regulation by Egr1 after overexpression and across the oestrous cycle. The Venn diagram from **Figure 4d** showing the overlap between genes annotated to female Egr1 gained-open regions and differentially expressed genes (DEGs) in ventral hippocampal neurons over the oestrous cycle (left). Taking the overlapping genes from this Venn diagram, an additional Venn diagram depicts that a subset of these genes also undergo chromatin accessibility changes across the oestrous cycle. Pro, proestrus; Die, dioestrus; EC, oestrus cycle.

**Extended Data Figure 10.**
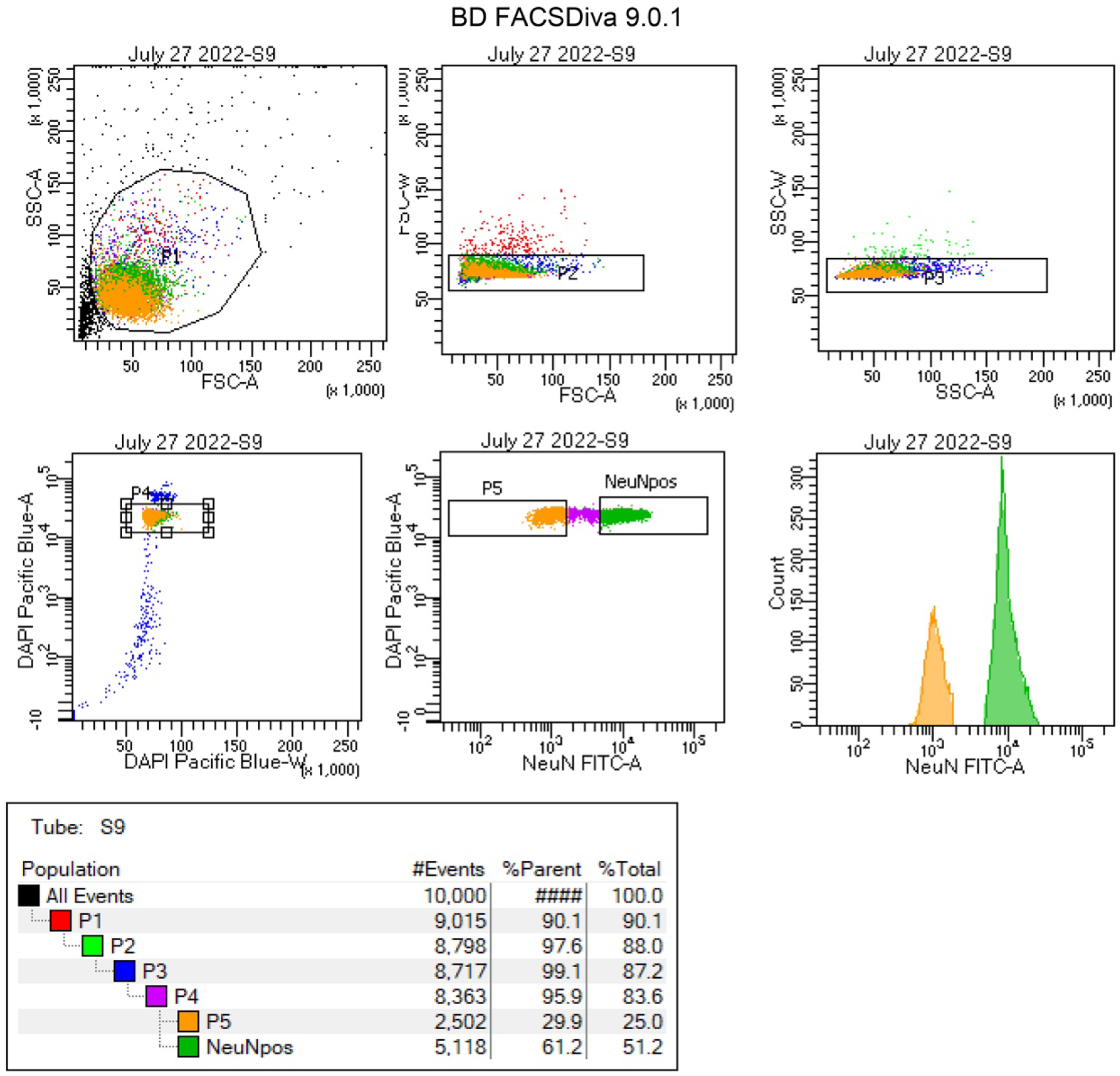
Isolation of neuronal nuclei using fluorescence-activated nuclei sorting (FANS) A representative FANS report which demonstrates the gating strategy used to: 1) separate the nuclei populations from debris (gates P1-P3); 2) sort only single nuclei, based on the DAPI fluorescent signal (gate P4); and, lastly, 3) separate NeuN+ (neuronal) nuclei (gate P6) from NeuN-(non-neuronal) nuclei (gate P5).

**Extended Data Figure 11.**
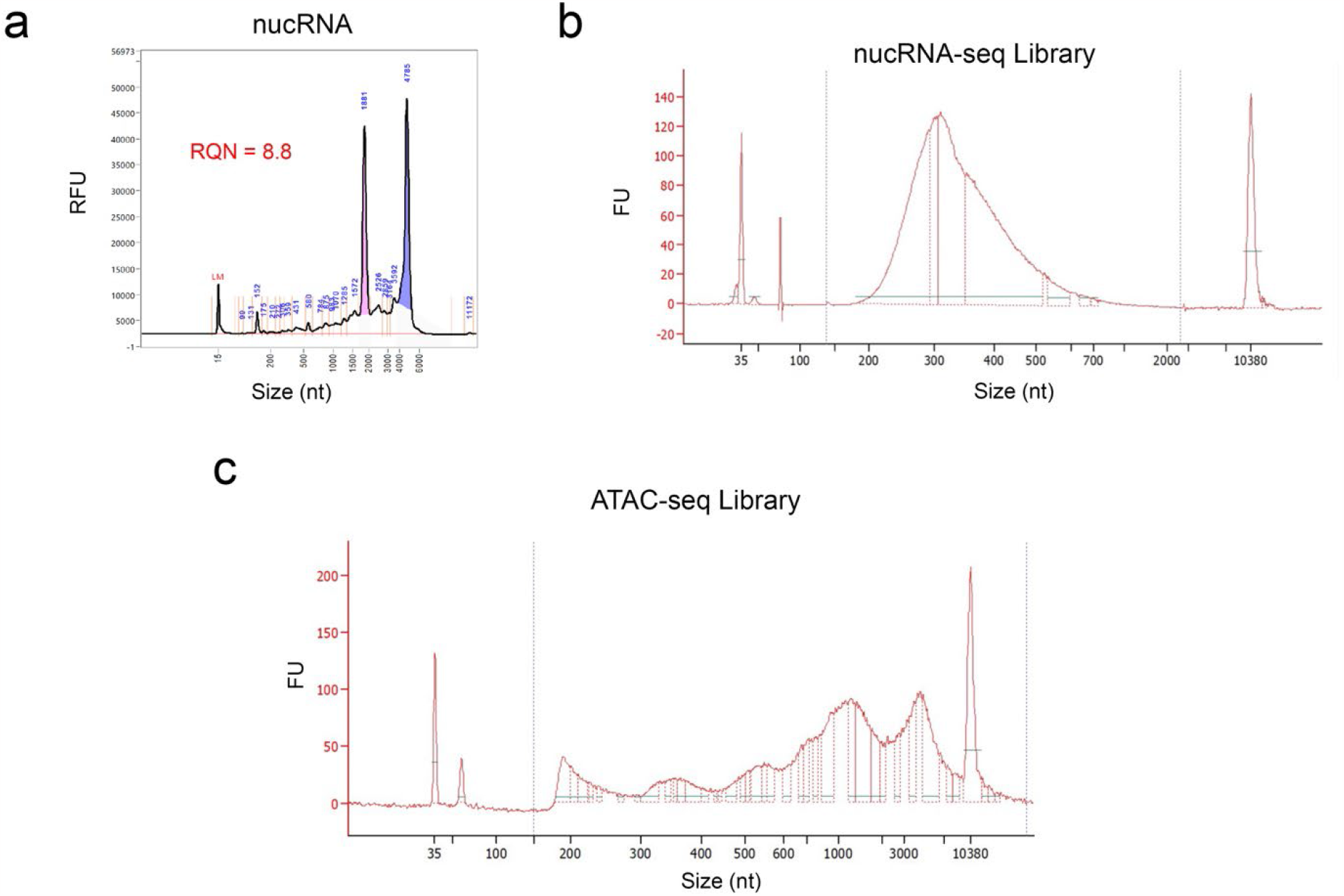
Quality control for nucRNA and sequencing libraries. **a**. A characteristic Fragment Analyzer trace of nuclear RNA (nucRNA) sample used in this study, with an RQN > 7 demonstrating high-quality RNA. Bioanalyzer traces of a cDNA library prepared from nucRNA (**b**) and from an ATAC-seq library (**c**) used in this study, with the ATAC-seq trace exhibiting a characteristic pattern of nucleosomal periodicity. RFU, relative fluorescence units; FU, fluorescence units.

